# ClipKIT: a multiple sequence alignment-trimming algorithm for accurate phylogenomic inference

**DOI:** 10.1101/2020.06.08.140384

**Authors:** Jacob L. Steenwyk, Thomas J. Buida, Yuanning Li, Xing-Xing Shen, Antonis Rokas

## Abstract

Highly divergent sites in multiple sequence alignments, which stem from erroneous inference of homology and saturation of substitutions, are thought to negatively impact phylogenetic inference. Trimming methods aim to remove these sites before phylogenetic inference, but recent analysis suggests that doing so can worsen inference. We introduce ClipKIT, a trimming method that instead aims to retain phylogenetically-informative sites; phylogenetic inference using ClipKIT-trimmed alignments is accurate, robust, and time-saving.

## Main

Multiple sequence alignment (MSA) of a set of homologous sequences is an essential step of molecular phylogenetics, the science of inferring evolutionary relationships from molecular sequence data. Errors in phylogenetic analysis can be caused by erroneously inferring site homology or saturation of multiple substitutions^1^, which often present as highly divergent sites. To address this issue, several methods “trim” or filter highly divergent sites using calculations of site/region dissimilarity from MSAs^1–3^. A beneficial by-product of MSA trimming, especially for studies that analyse hundreds of MSAs from thousands of taxa^4^, is that trimming MSAs reduces the computational time and memory required for phylogenomic inference. Nowadays, MSA trimming is a routine part of molecular phylogenetic inference^5^.

Despite the overwhelming success of MSA trimming methods, a recent analysis by Tan *et al*. revealed that trimming often decreases, rather than increases, accuracy of phylogenetic inference^6^. This decrease suggests that current methods may remove phylogenetically-informative sites (e.g., parsimony-informative and variable sites) that have previously been shown to contribute to phylogenetic accuracy^7^. Furthermore, Tan *et al*. showed that phylogenetic inaccuracy is positively associated with the number of removed sites^6^, revealing a speed-accuracy trade-off wherein trimmed MSAs decrease the computation time of phylogenetic inference but at the cost of reduced accuracy. More broadly, these findings highlight the need for alternative MSA trimming strategies.

To address this need, we developed ClipKIT, an MSA-trimming algorithm based on a conceptually novel framework. Rather than aiming to identify highly divergent sites/regions in MSAs, ClipKIT instead focuses on identifying and retaining phylogenetically-informative sites. Specifically, ClipKIT has five trimming modes:

1. kpi: retain only parsimony-informative sites (i.e., sites with at least two characters that each occur at least twice), which are associated with phylogenetic signal^7^,
2. kpic: retain both parsimony-informative and constant sites, the latter of which helps inform parameter estimation in substitution models^8^,
3. gappy: trimming based on site gappyness (i.e., sites with ≤90% gaps are kept),
4. kpi-gappy: mode 1 combined with mode 3, and
5. kpic-gappy: mode 2 combined with mode 3.

To test the efficacy of ClipKIT, we examined the accuracy and support of single-gene and species-level phylogenetic trees inferred from untrimmed MSAs and MSAs trimmed using 13 different approaches (Table 1) across four empirical genome-scale datasets and four simulated datasets. The four empirical datasets correspond to the untrimmed amino acid and nucleotide MSAs from 24 mammals (N_alignments_=4,004) and 12 budding yeasts (N_alignments_=5,664)^7^. The four simulated datasets (N_alignments_=50 alignments per dataset or 200 total) stem from simulated nucleotide sequence evolution along the species phylogeny of 93 filamentous fungi^9^, and from simulated amino acid sequence evolution along the species phylogenies of 70 metazoans^10^, 46 flowering plants^11^, and 96 budding yeasts^12^. Simulated sequences were generated with INDELible, v1.03^13^. MSAs were trimmed using popular alignment trimming software (Table 1) generating a total of 138,152 MSAs [(4,004 mammalian + 5,664 yeast + 200 simulated MSAs) * 14 treatments = 138,152 MSAs]. However, Gblocks and BMGE with an entropy threshold of 0.3 were not used for performance assessment of simulated datasets because they frequently removed entire MSAs.

**Table 1.**
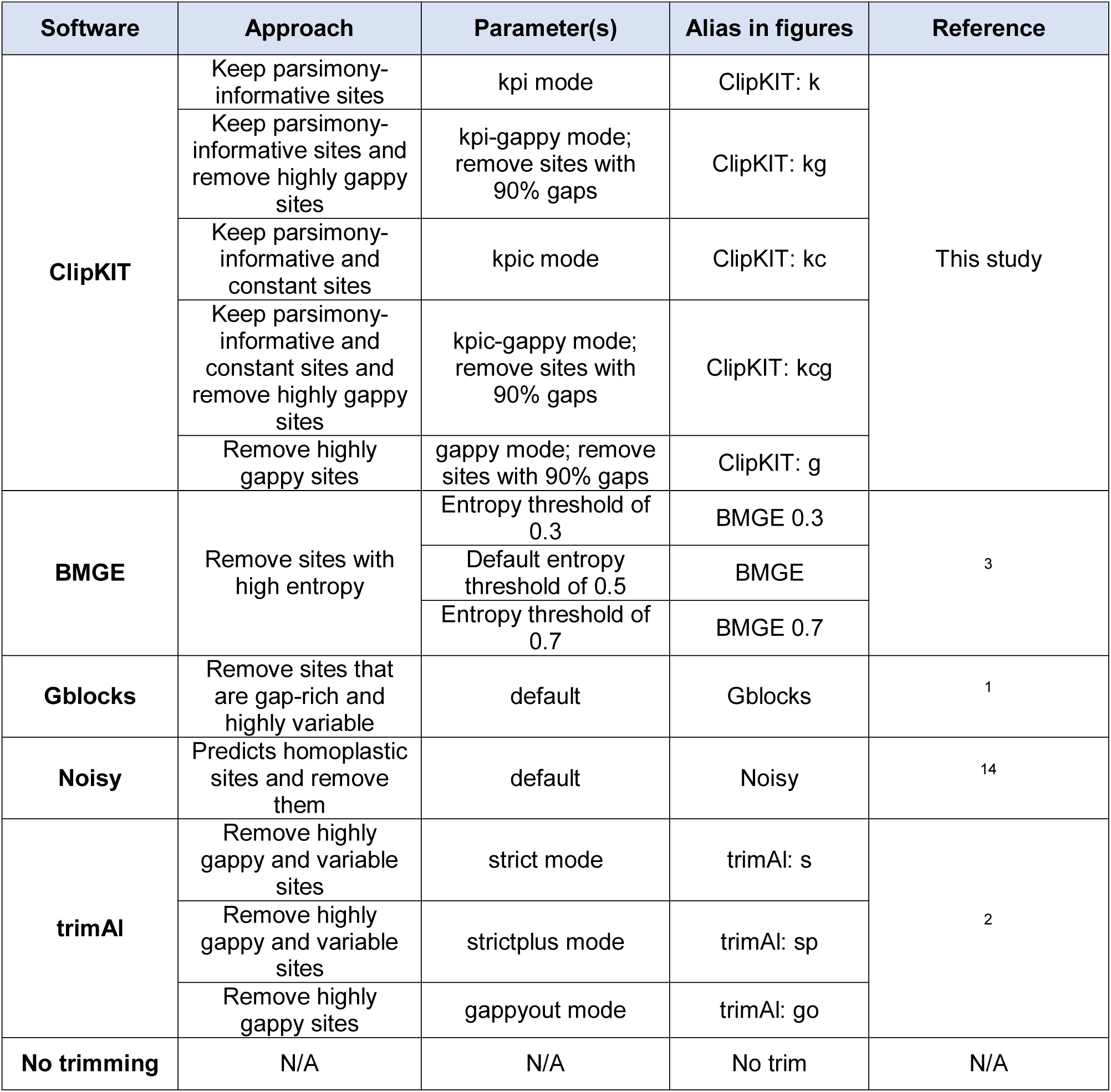
Multiple sequence alignment (MSA)-trimming methods tested. Each MSA-trimming strategy tested by our analysis, a general description of its trimming approach, and parameters are described here.

We next conducted single-gene phylogenetic inference with all MSAs using IQ-TREE, v1.6.11^8^. Tree accuracy was measured using normalized Robinson-Foulds (nRF) distances as calculated by ape, v5.3^15^, R package (https://cran.r-project.org/), by comparing the inferred gene phylogenies to their species phylogenies. Tree support was measured using average bipartition support (ABS) from 5,000 ultrafast bootstrap approximations in IQ-TREE. To determine if alignment trimmers resulted in substantially different alignment lengths, nRF values, and ABS values, we conducted principal component analysis.

We found that the 14 approaches examined occupied distinct regions of feature space suggestive of substantial differences between MSAs (Figure 1). Variation in feature space was largely driven by nRF and ABS measures along the first dimension and alignment length along the second dimension for both empirical and simulated datasets (Figure S1). In empirical datasets, we found that some ClipKIT modes removed few sites while others removed many and, at times, the most sites (Supplementary figure 2). Among simulated datasets, ClipKIT trimmed substantial portions of MSAs but variation was observed across MSAs and datasets (Supplementary figure 3). Examination of nRF and ABS values revealed ClipKIT performed well, and at times the best, among the MSA-trimming approaches tested, suggesting that phylogenetic inferences made with ClipKIT-trimmed MSAs were both accurate and well supported (Supplementary figure 4 and 5). Finally, counter to previous evidence suggestive of a trade-off between trimming and phylogenetic accuracy^6^, we found that ClipKIT aggressively trimmed MSAs in the empirical datasets without compromising phylogenetic tree accuracy and support.

**Figure 1.**
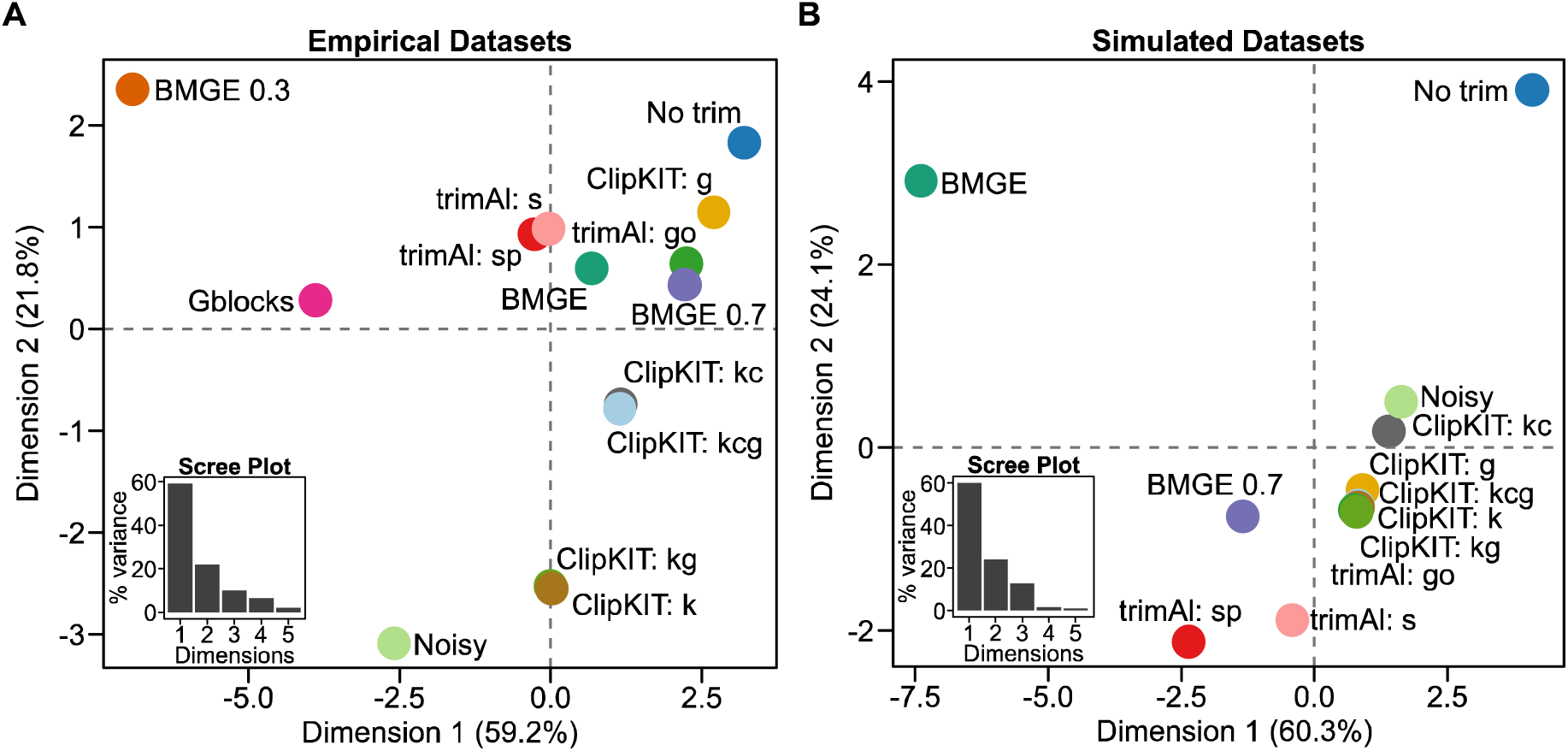
Alignment trimming algorithms differ in resulting multiple sequence alignments and metrics of phylogenetic tree accuracy and support. Principal component analysis of alignment length, nRF, and ABS values across various MSA trimming approaches for four empirical datasets (A) and four simulated datasets (B). Insets of scree plots depict the percentage of variation explained (y-axis) for the first five dimensions (x-axis). Data was scaled prior to conducting principal component analysis.

To obtain a summary of overall performance, we ranked the 14 approaches’ performance for each dataset using objective desirability-based integration of nRF and ABS values^16^ (Figure 2). We found that the five ClipKIT modes outperformed all other alignment trimming software for amino acid sequences in the empirical mammalian dataset (Figure 2A) as well as the simulated datasets of metazoan and flowering plant sequences (Figure 2E and F). Other software that performed well included trimAl with the ‘gappyout’ parameter for empirical datasets and Noisy for simulated datasets^2,14^. To evaluate MSA trimming algorithm performance for empirical and simulated datasets, we examined average ranks across each set of four datasets and found ClipKIT modes were among the best performing (Figure 2 I-J). Specifically, among empirical sequences, ClipKIT’s gappy mode outperformed all other approaches followed by no trimming, trimAl with the ‘gappyout’ parameter, and then four other ClipKIT modes (Figure 2I); among simulated sequences, no trimming ranked best followed by all five ClipKIT modes (Figure 2J). These results suggest that ClipKIT, which focuses on retaining phylogenetically-informative sites, was on par with no trimming and frequently outperformed approaches that focus on removing highly divergent sites.

**Figure 2.**
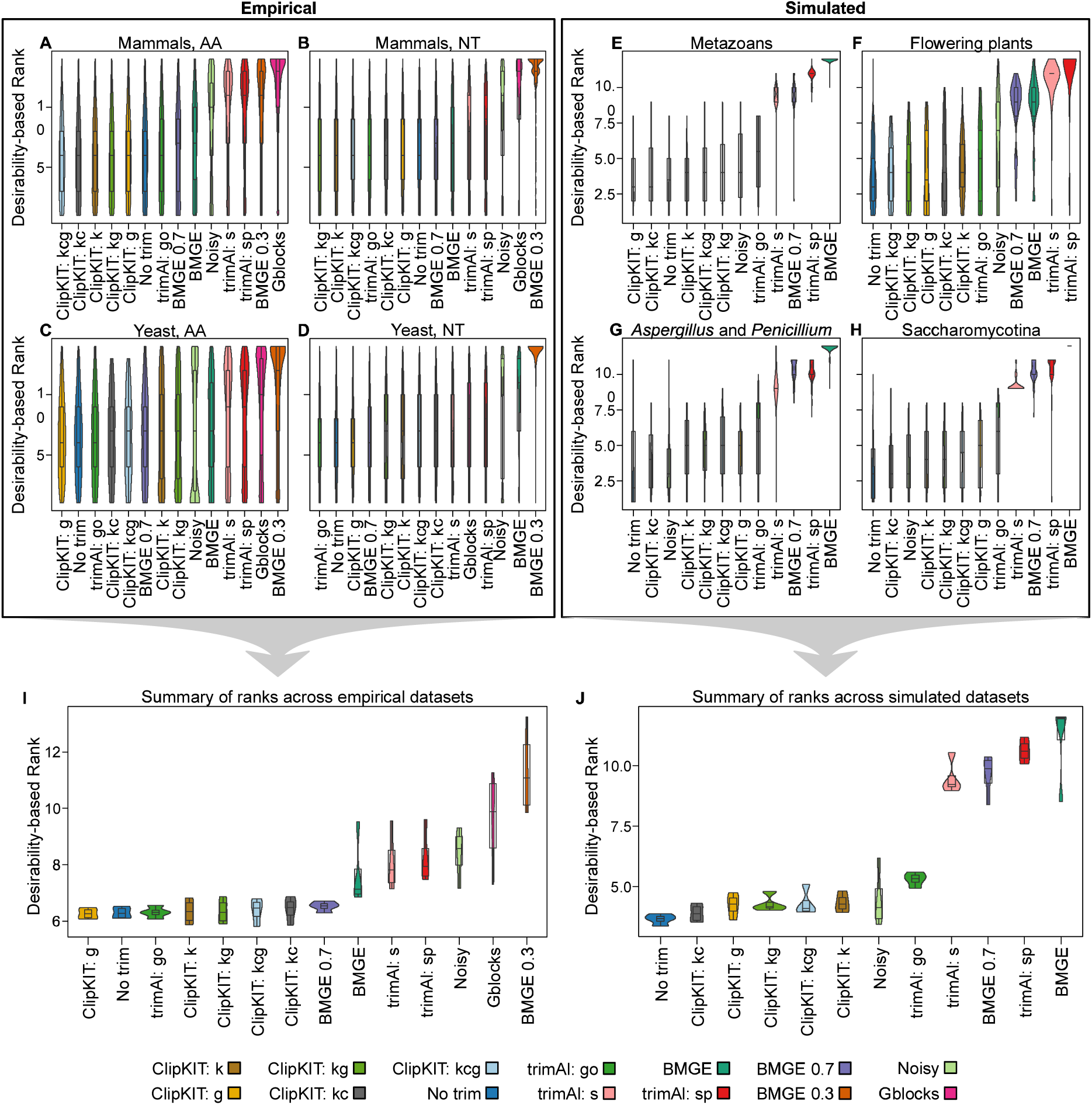
ClipKIT is a top-performing software for trimming multiple sequence alignments. Desirability-based integration of accuracy and support metrics per MSA facilitated the comparison of relative software performance for empirical (A-D) and simulated (E-H) datasets. Examination of software performance for individual datasets and average performance across empirical (I) and simulated (J) datasets revealed ClipKIT is a top-performing software. MSA trimming approaches are ordered along the x-axis from the highest-performing software to the lowest-performing software according to average desirability-based rank. Boxplots embedded in violin plots have upper, middle, and lower hinges that represent the first, second, and third quartiles. Whiskers extend to 1.5 times the interquartile range.

To evaluate the performance of the 14 approaches, including ClipKIT-based ones, for species-level phylogenetic inference, we conducted concatenation and coalescence-based phylogenetic inference using IQ-TREE and Astral, v 5.7.3^17^, respectively. We found that all MSA-trimming software resulted in nearly identical and well supported phylogenies (Supplementary figures 6-8). Among simulated datasets, we found that ClipKIT approaches reduced computation time by an average of ∼20% compared to no trimming.

In summary, ClipKIT performed consistently well across empirical and simulated data. These results suggest that MSA-trimming focused on retaining phylogenetically-informative sites often outperformed approaches focused solely on removing highly divergent sites and had similar performance to no trimming (but significantly reduced computation time). We anticipate ClipKIT will be useful for phylogenomic inference and the quest to build the tree of life.

## Methods

ClipKIT is a standalone software written in the Python programming language (https://www.python.org/) and is available from GitHub, https://github.com/JLSteenwyk/ClipKIT, and PyPi, https://pypi.org/. ClipKIT differs from most multiple sequence alignment (MSA) trimming methods because it focuses on identifying and retaining phylogenetically-informative sites from MSAs rather than removing highly divergent ones. To do so, ClipKIT conducts site-by-site examination of MSAs and determines whether they should be retained or trimmed based on the mode of ClipKIT being used. ClipKIT has five trimming modes:

1. kpi: a mode that retains sites that are parsimony-informative, which is specified with the following command:

~~~
clipkit <MSA> -m kpi;
~~~
2. kpic: a mode that retains sites that are either parsimony-informative or constant, which is specified with the following command:

~~~
clipkit <MSA> -m kpic;
~~~
3. gappy: a mode that retains sites that are not gappy-rich (defined as sites with ≤90% gaps), which is specified with the following command:

~~~
clipkit <MSA> -m gappy,
~~~

alternatively, gappy-based trimming is the default mode and the same style of trimming can be achieved with the following command:

~~~
clipkit <MSA>;
~~~
4. kpi-gappy: a combination of mode 1 and mode 3, which is specified with the following command:

~~~
clipkit <MSA> -m kpi-gappy;
~~~ and
5. kpic-gappy: a combination of mode 2 and mode 3, which is specified with the following command:

~~~
clipkit <MSA> -m kpic-gappy.
~~~

All output files have the same name as the input files with the addition of the suffix “.clipkit.” Users can specify output files names with the -o/--output option. For example, an alignment may have the output name “ClipKIT_trimmed_aln.fa” with the following command:

~~~
clipkit <MSA> -o ClipKIT_trimmed_aln.fa.
~~~

To enable users to fine-tune alignment trimming parameters, we provide an additional option for users to specify their own gappyness threshold, which can range between zero and one. For example, to retain sites with ≤95% gaps, the following command would be used:

~~~
clipkit <MSA> -g 0.95
~~~

In practice, we recommend the gaps parameter never be set too low because trimming may remove too many sites, which may lead to worse phylogenetic inferences^7^.

To enable users to examine the trimmed sites/regions from MSAs, we have also implemented a logging option in ClipKIT. When used, the logging option outputs an additional four-column file with the following information: column 1, position in the alignment (starting at 1); column 2, whether or not the site was trimmed or kept; column 3, reports if the site was parsimony-informative, constant, or neither and; column 4, reports the gappyness of the site. Log files are generated using the -l/--log option:

~~~
clipkit <MSA> -l
~~~

We anticipate this information will be helpful for alignment diagnostics, fine-tuning of trimming parameters, and other reasons.

To enable seamless integration of ClipKIT into pre-existing pipelines, eight file types can be used as input. More specifically, ClipKIT can input and output *fasta, clustal, maf, mauve, phylip, phylip-sequential, phylip-relaxed*, and *stockholm* formatted MSAs. By default, ClipKIT automatically determines the input file format and creates an output file of the same format; however, users can specify either with the -if/--input_file_format and -of/--output_file_format options. For example, an input file of *fasta* format and a desired output file of *clustal* format can be specified using the following command:

~~~
clipkit <MSA> -if fasta -of clustal
~~~

Recent analyses indicate that ∼28% of available computational tools fail to install due to implementation errors^18^. To overcome this hurdle and ensure archival stability of ClipKIT, we implemented state-of-the-art software development practices and design principles. More specifically, ClipKIT is composed of highly modular, extensible, and reusable code, which allows for easy debugging and seamless integration of new functions and features. We wrote a total of 118 unit and integration tests resulting in 97% code coverage. This high level of coverage was achieved due to ClipKIT’s exemplary engineering practices. We also implemented a robust continuous integration (CI) pipeline to automatically build, package, and test ClipKIT whenever code is modified. This CI pipeline runs a testing matrix for Python versions 3.6, 3.7, and 3.8. Given the current configuration, building and testing ClipKIT for future versions of Python will be trivial to implement. Lastly, central ClipKIT functions rely on few dependencies (i.e., BioPython^19^ and NumPy^20^). In summary, we have taken several measures to ensure ClipKIT implements a method that trims MSAs without sacrificing accuracy of phylogenetic inference as well as ensures that it will be a long-lasting computational tool for the field of molecular phylogenetics.

### Practical considerations

Although ClipKIT performed well across empirical genome-scale and simulated datasets, we acknowledge that testing every possible evolutionary scenario is impossible. This is further complicated by the lack of large-scale phylogenomic data matrices in which the true evolutionary relationships among organisms are known. Therefore, we recommend using multiple trimming modes available in ClipKIT and examining the resulting ABS values for trees. Considering high ABS values often corresponded to lower nRF values (Supplementary Figure 4 and 5), using the resulting phylogeny with the highest ABS value may be representative of the phylogeny that most closely resembles the sequences’ true evolutionary history. This may require substantially greater computation time. To potentially ameliorate the computation time issue that may arise, we recommend creating subsets of larger datasets that span alignments of various lengths and testing multiple trimming modes on the reduced dataset.

Although constant sites are thought to be important for informing parameters of substitution models^8^, we observed variation in the performance of ClipKIT modes that retain only parsimony-informative sites (modes: kpi and kpi-gappy) and modes that retain parsimony-informative and constant sties (modes: kpic and kpic-gappy). More specifically, at times modes kpi and kpi-gappy outperformed kpic and kpic-gappy and vice versa suggesting constant sites may not be as informative to substitution models. However, we note that trimming nucleotide sequences with modes kpi and kpi-gappy may warrant ascertainment bias correction for nucleotide sequences because constant sites are absent from the trimmed alignments.

## Software availability

ClipKIT is available from GitHub, https://github.com/JLSteenwyk/ClipKIT, and PyPi, https://pypi.org/project/clipkit.

## Data availability

All alignments and phylogenies inferred in this study will be available from figshare (doi: 10.6084/m9.figshare.12401618) upon publication.

