## Supplementary Tables and Figures for "ClipKIT: a multiple sequence alignment-trimming algorithm for accurate phylogenomic inference"

[2] 9 City Place #312, Nashville, TN 37209, United States of America

[3] Institute of Insect Sciences, Ministry of Agriculture Key Lab of Molecular Biology of Crop Pathogens and Insects, College of Agriculture and Biotechnology, Zhejiang University, Hangzhou 310058, China

#### Supplementary methods

To determine how well ClipKIT-trimmed MSAs performed during phylogenetic inference, we used empirical and simulated datasets. Information on how the empirical datasets were obtained are in the main text. Simulated alignments were generated using INDELible, v1.03[1]. Nucleotide alignments for filamentous fungi were generated using the general time reversible (GTR) substitution model[2]. Additional parameters specified were state frequencies values of 0.1, 0.2, 0.3, and 0.4 for T, C, A, and G nucleotides, respectively. We specified the substitution rate matrix using the scheme outlined in Supplementary table 1.

| | T | C | A | G |
|---|---|---|---|---|
| **T** | - | 0.2 | 0.4 | 0.6 |
| **C** | 0.2 | - | 0.8 | 1.2 |
| **A** | 0.4 | 0.8 | - | 1 |
| **G** | 0.6 | 1.2 | 1 | - |

**Supplementary table 1. GTR substitution rate matrix.** Simulated nucleotide alignments were generated using this substitution rate matrix.

Insertion and deletion rates were set to be 5% as frequent as single substitutions. Insertion and deletions occurred according to the power law distribution (a=1.7, M=500). The tree's root length was set to 1,000. For amino acids, all parameters were the same except the insertion and deletion rates were set to be 1% as frequent as single substitutions using the WAG model of substitutions, which was also used to specify state frequencies[3].

We use IQ-TREE[4] for maximum likelihood phylogenetic inference. For nucleotide sequences, we used a GTR substitution model[5] with empirical base frequencies and a discrete Gamma model with four rate categories[6] or "GTR+F+G"; for amino acid sequences, we used the general WAG model of substitutions[3] with empirical base frequencies and a discrete Gamma model with four rate categories[6] or "WAG+F+G."

### Supplementary figures and legends

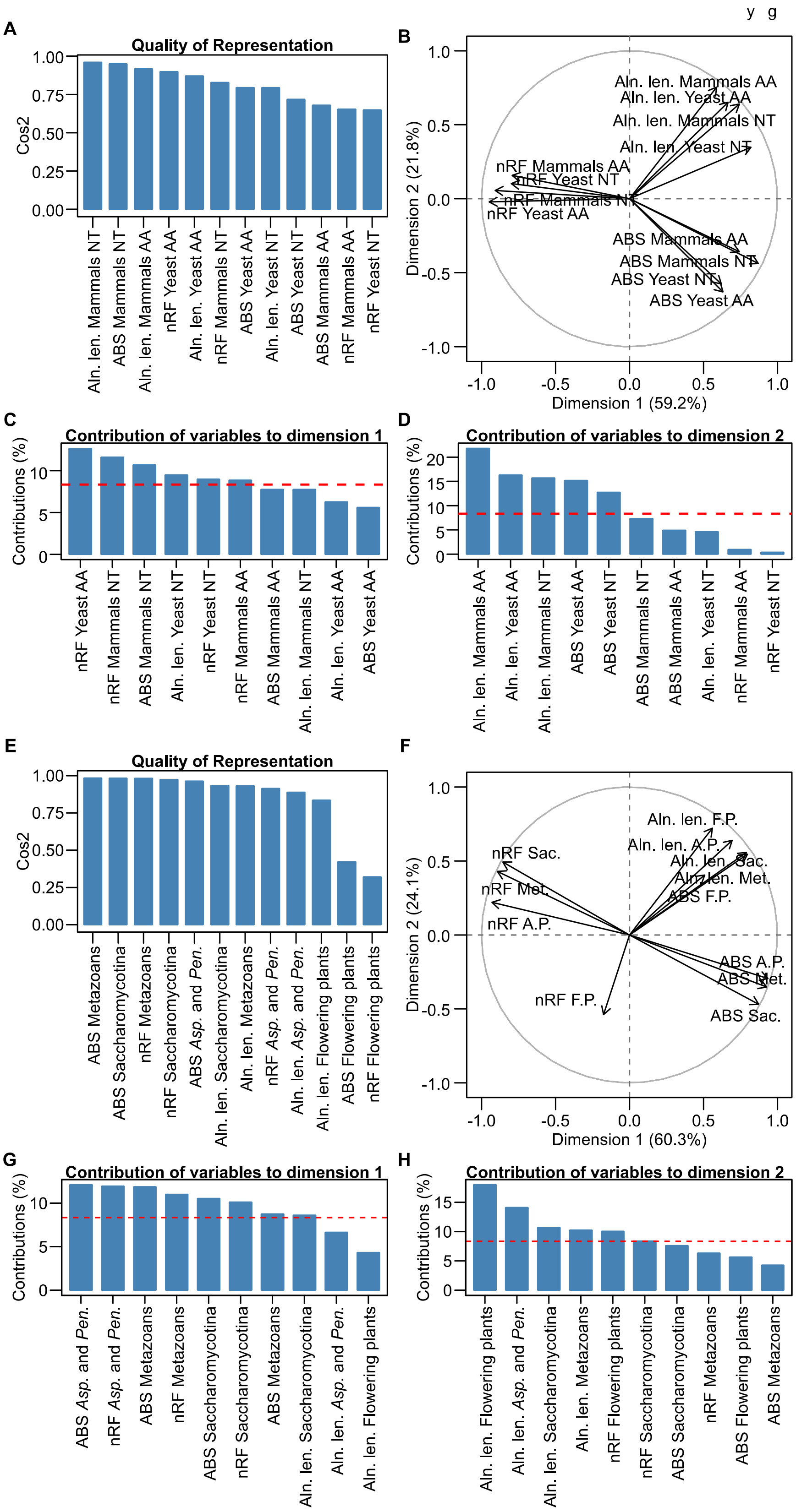

**Supplementary figure 1. Data are well represented in principal component analysis.** Examination of the factors contributing how all trimming approaches varied in feature space for empirical datasets are represented in A-D and simulated datasets are represented in E-H. Examination of variable representation along the first and second dimensions of the principal component analysis (see Figure 1) revealed alignment length, nRF, and ABS were well represented in empirical (A) and simulated (E) datasets. Variable correlation plots for empirical (B) and simulated (F) datasets show the relationship among variables. Broadly, we found that variable types (i.e., alignment length, nRF, and ABS) were correlated with one another across datasets. Examination of contribution of variables along the first and second dimensions for empirical datasets (C and D, respectively) as well as simulated datasets (G and H, respectively) revealed that ABS and nRF contributed the most along the first dimension and alignment length contributed the most along the second dimension. In these figures, the red dashed line represents the expected average contribution if all variables contributed equally. Abbreviations used in the figure are as follows: alignment length: Aln. len.; Saccharomycotina: Sac.; Metazoans: Met.; Flowering plants: F.P.; *Aspergillus* and *Penicillium*: A.P.

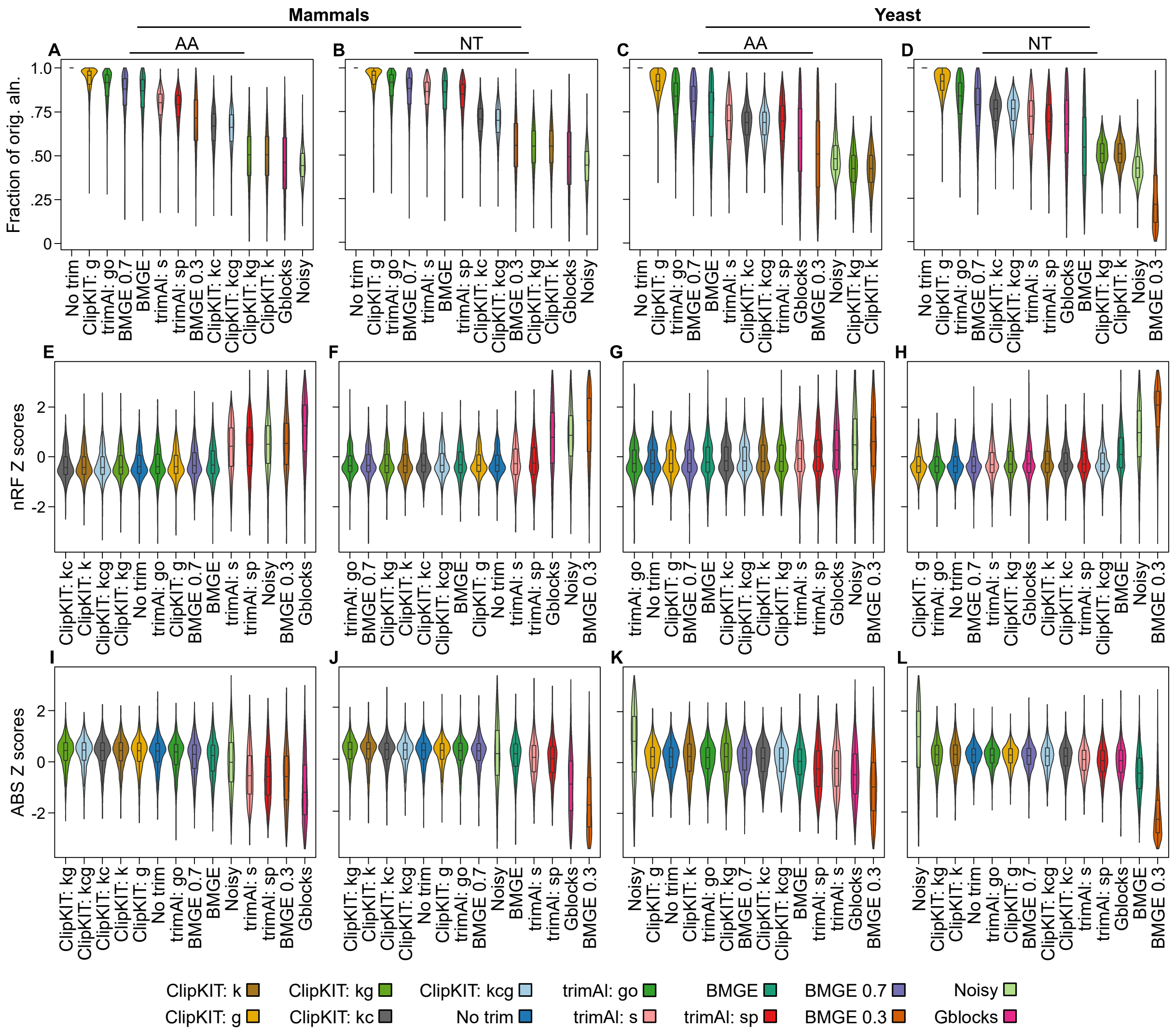


**Supplementary figure 2. Lengths of trimmed MSAs and the associated nRF and ABS values across empirical datasets.** For the mammalian amino acid (AA) and nucleotide (NT) sequences (A and B) as well as the yeast AA and NT sequences (C and D), wide variation was observed in the fraction of the original alignment trimmed by different trimming approaches. MSA trimming approaches are ordered along the x-axis from the least aggressive trimmer to the most aggressive trimmer according to the average fraction of the original alignment in that remained in the trimmed alignment (y-axis). Distributions of nRF (E-H) and ABS (I-L) values were subsequently examined. MSA trimming approaches are ordered along the x-axis from the highest-performing software to the lowest-performing software according to average nRF or

ABS value. nRF and ABS values were Z transformed per gene prior to plotting their distribution. Boxplots embedded in violin plots have upper, middle, and lower hinges that represent the first, second, and third quartiles. Whiskers extend to 1.5 times the interquartile range. Note that the yeast dataset (N=12) is at the lower threshold of Noisy's recommended minimum number of sequences[7].

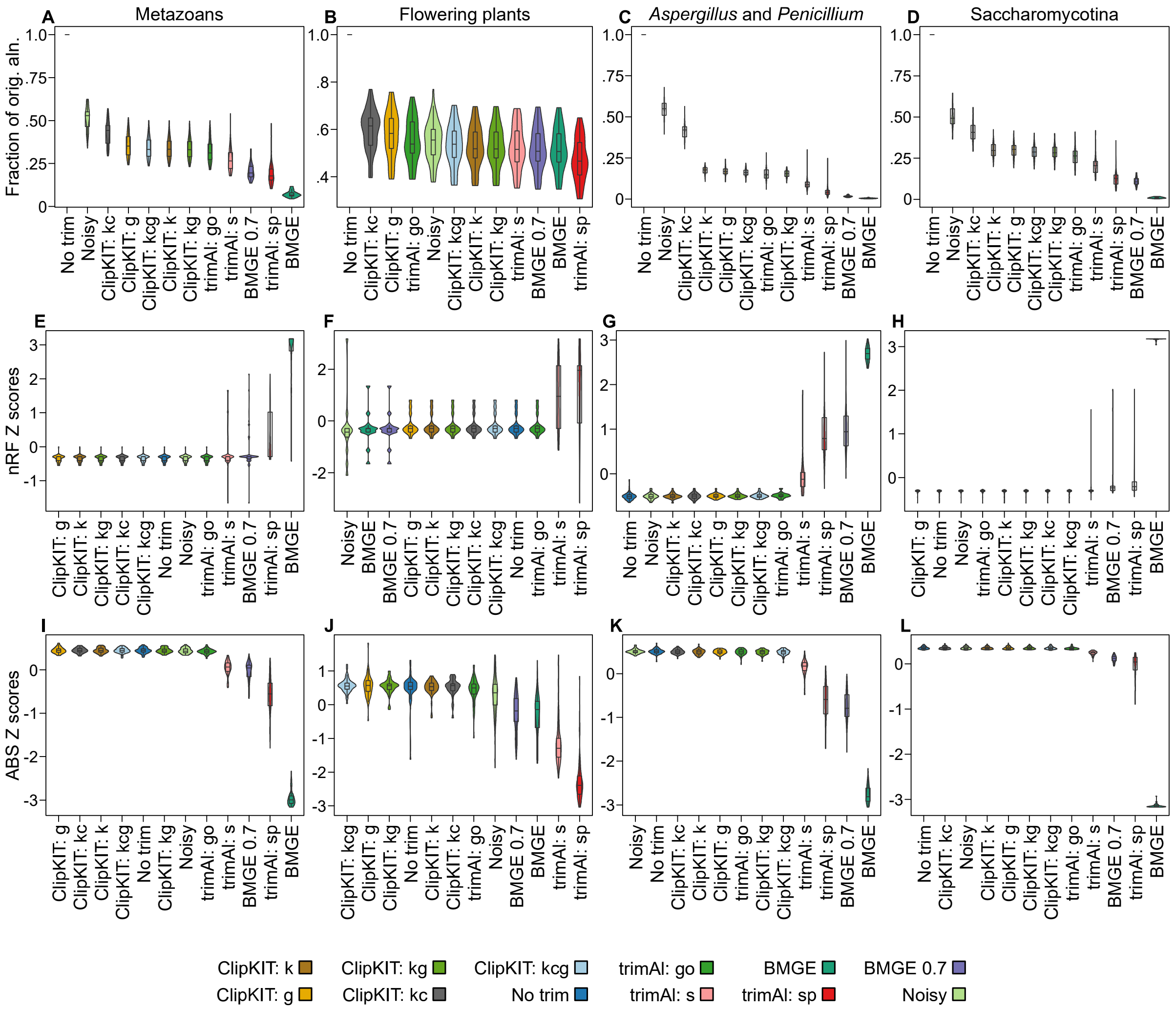


**Supplementary figure 3. Lengths of trimmed MSAs and the associated nRF and ABS values across simulated datasets.** Extensive variation was observed in the fraction of the original alignment trimmed by the various MSA trimming approaches (A-D). MSA trimming approaches are ordered along the x-axis from the least aggressive trimmer to the most aggressive trimmer according to the average fraction of the original alignment that remained in the trimmed alignment (y-axis). Among the various datasets, we examined the distributions of nRF (E-H) and ABS (I-L) values. MSA trimming approaches are ordered along the x-axis from the highest-performing software to the lowest-performing software according to average nRF or ABS value. Prior to plotting, nRF and ABS values were Z transformed on a per gene basis. Boxplots embedded in violin plots have upper, middle, and lower hinges that represent the first, second, and third quartiles. Whiskers extend to 1.5 times the interquartile range.

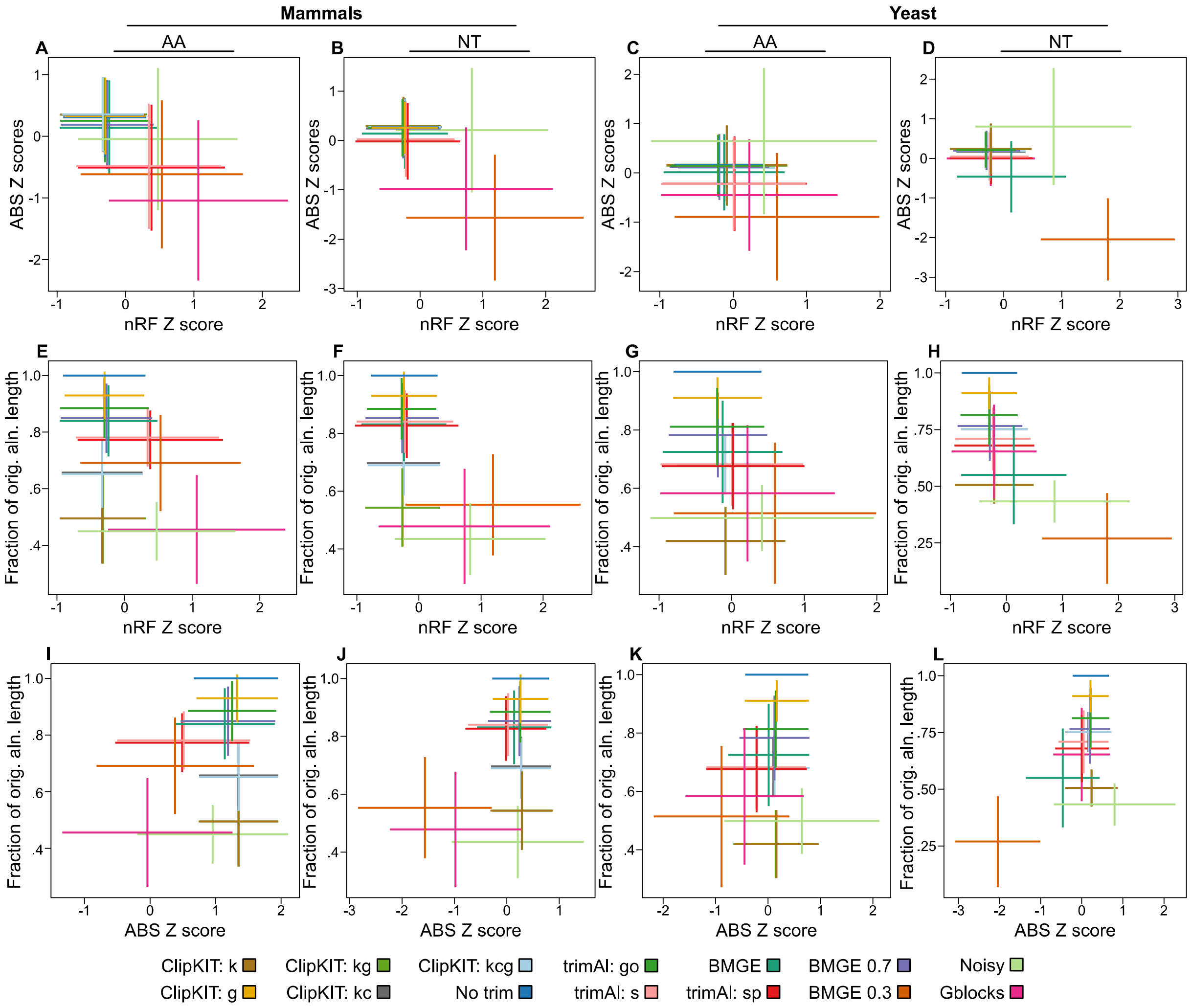


**Supplementary figure 4. Pairwise examination of the relationship between alignment length, nRF, and ABS values among empirical datasets.** Broadly, we found that higher ABS values corresponded with lower nRF values across the various datasets (A-D). Additionally, lower nRF values were typically associated with shorter alignment lengths (E-H). The only MSA trimmers that did not follow this trend were the ClipKIT modes kpic, kpic-gappy, kpi and kpi-gappy, where the alignments were shorter than others but resulted in accurate phylogenetic inferences. Similarly, we found longer alignments were associated with higher ABS values (I-L). Again, we found that the only alignment trimming algorithm that resulted in substantially shorter alignments, but which produced well supported phylogenetic treess, were the ClipKIT modes kpic, kpic-gappy, kpi and kpi-gappy. This suggests that ClipKIT was able to trim substantial portions of alignments without compromising phylogenetic accuracy and support. Bidirectional error bars extend one standard deviation from the mean. Error bars cross at the average of the two variables being examined.

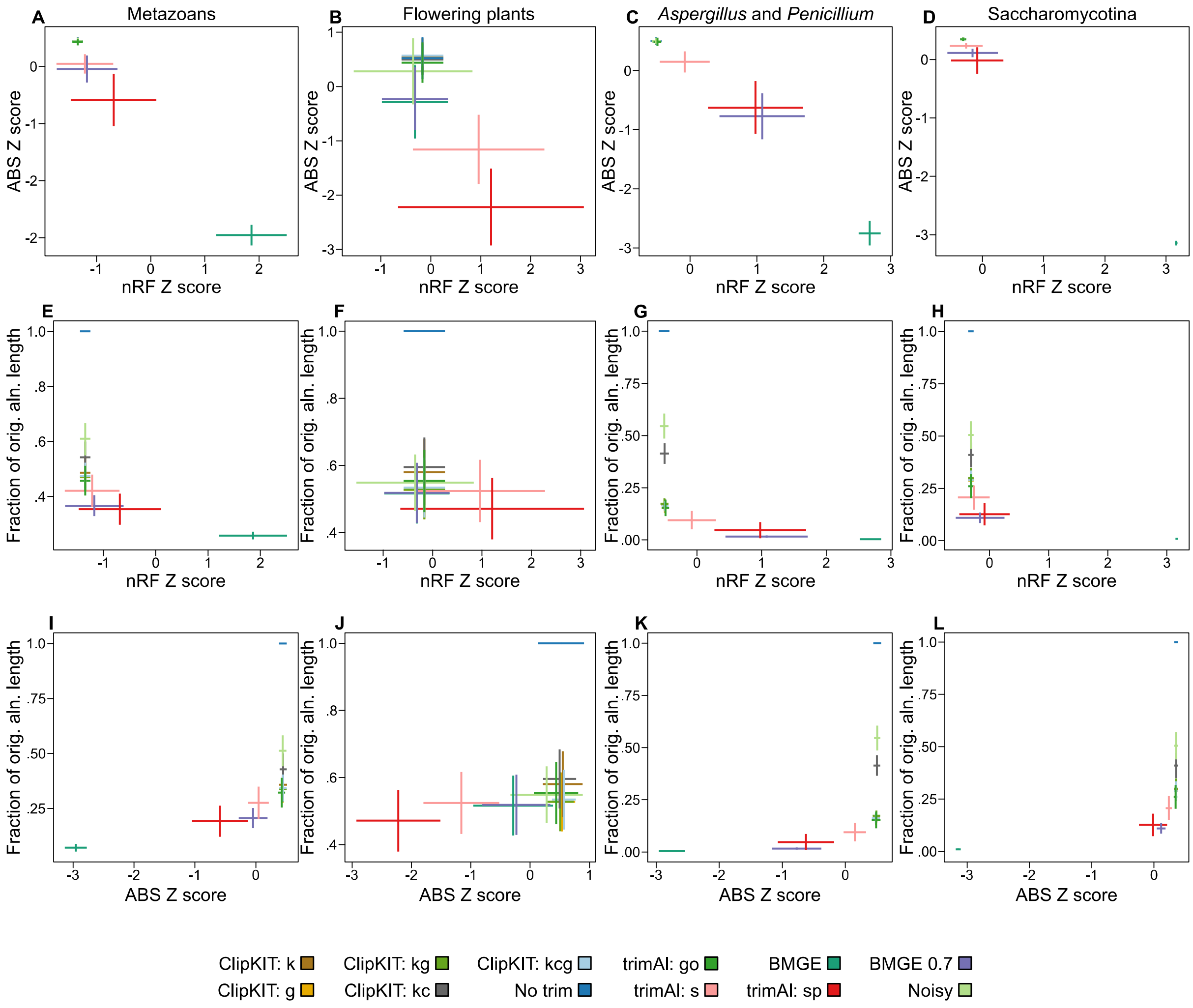


**Supplementary figure 5. Pairwise examination of the relationship between alignment length, nRF, and ABS values among simulated datasets.** Higher ABS values were often associated with lower nRF values (A-D). Lower nRF values were often associated with shorter alignment lengths (E-H). Longer alignments were often associated with higher ABS values (I-L). Bidirectional error bars extend one standard deviation from the mean. Error bars cross at the average of the two variables being examined.

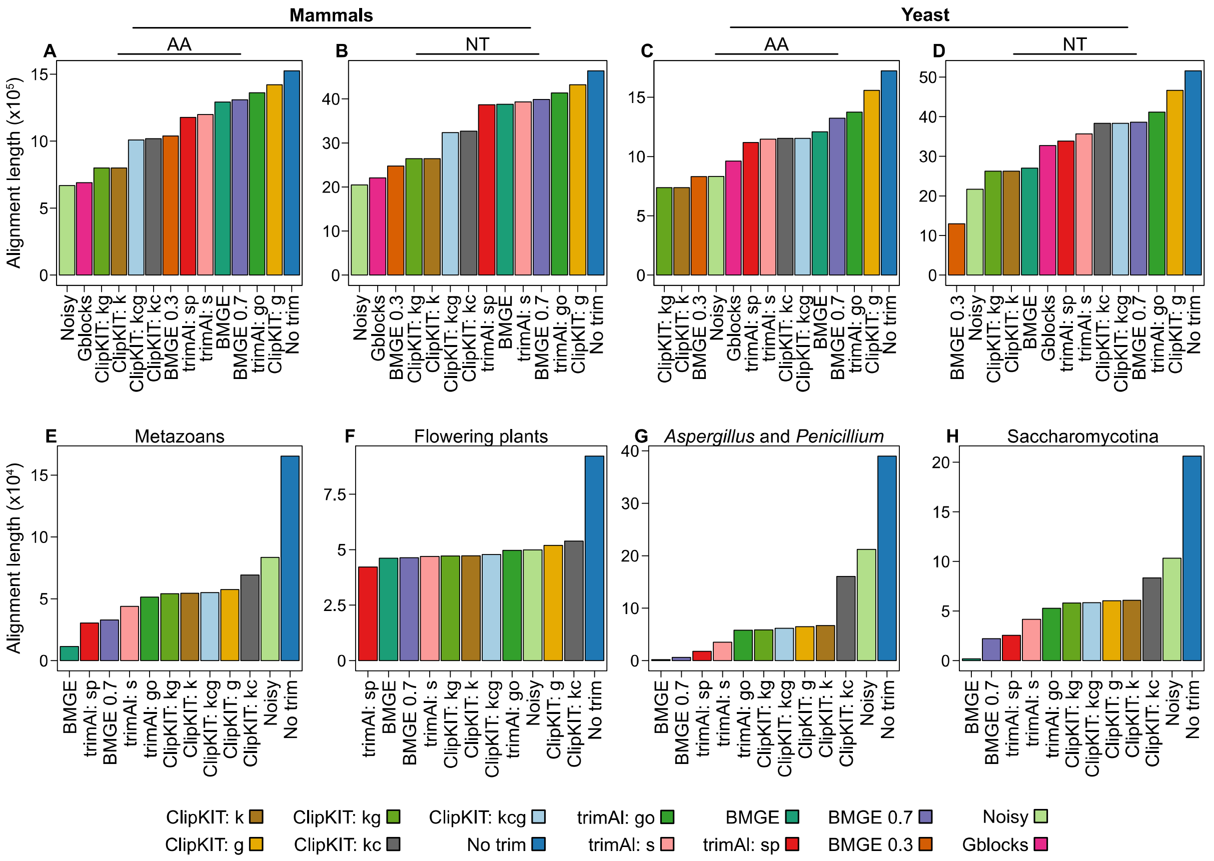


**Supplementary figure 6. Concatenated alignment lengths varied by dataset and alignment trimming method.** Alignment lengths of concatenated data matrices for mammals (A-B), yeasts (C-D), which are the empirical datasets, and metazoans (E), flowering plants (F), *Aspergillus* and *Penicillium* (G), and Saccharomycotina yeasts (H), which are simulated datasets, varied greatly. For example, ClipKIT with the gappy mode trimmed the least out of all methods among the empirical datasets while Noisy often trimmed the least among simulated datasets. MSA trimming approaches are ordered along the x-axis from the ones that trimmed the most to those that trimmed the least.

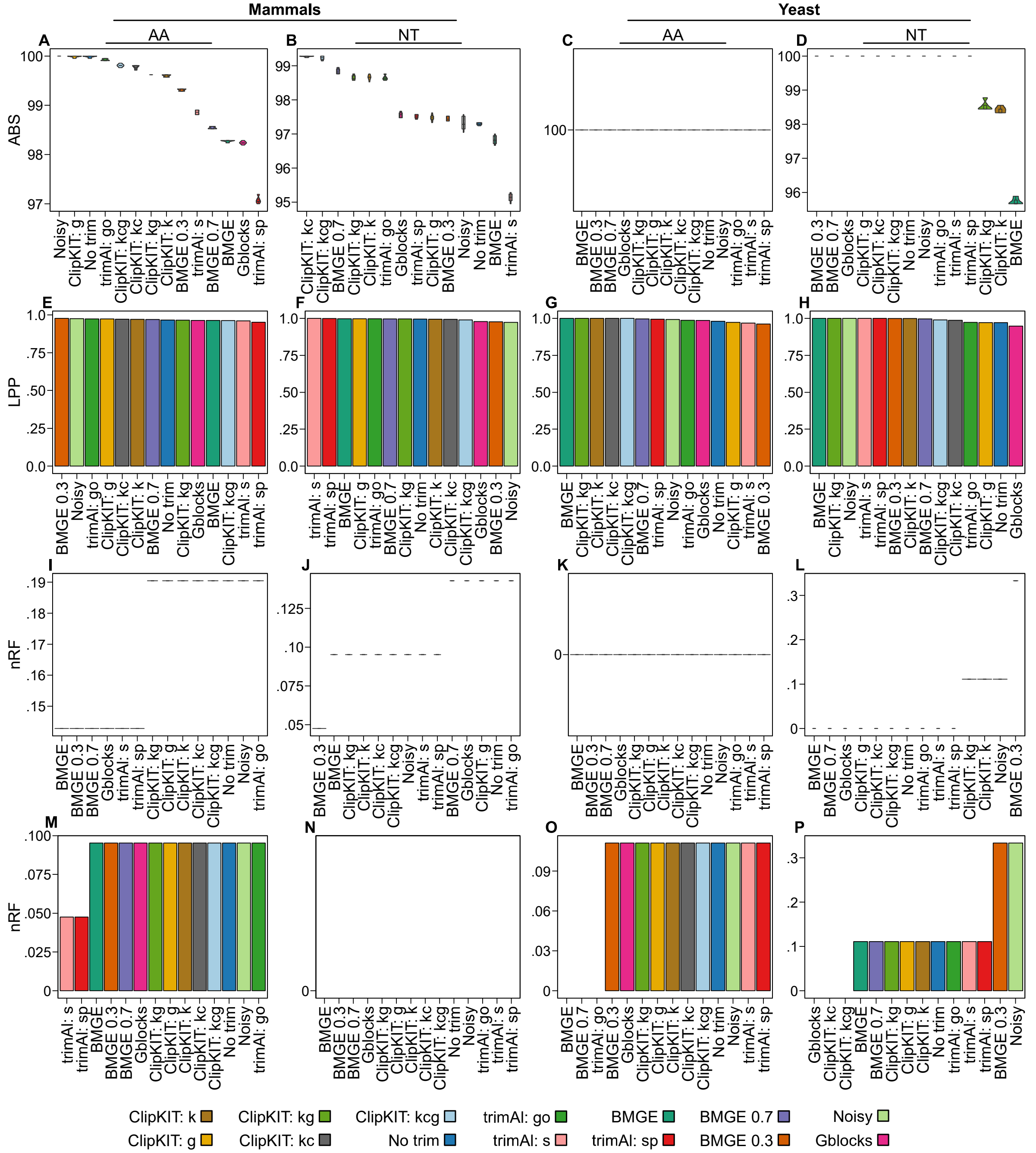


**Supplementary figure 7. Alignment trimming approaches resulted in nearly identical metrics of accuracy and support for species-level inferences using empirical datasets.** Species-level phylogenies were inferred using a concatenation and coalescence approaches. All phylogenies inferred received nearly full support using the concatenation (A-D) and coalescence approaches (E-H). Similarly, phylogenies inferred using the concatenation (I-L) and coalescence approaches (M-P) were accurate. Variation species-level inferences for these datasets has been previously reported[8]. Thus, we are considering these phylogenies nearly identifical. Distributions among concatenation-based metrics stem from five independent tree searches.

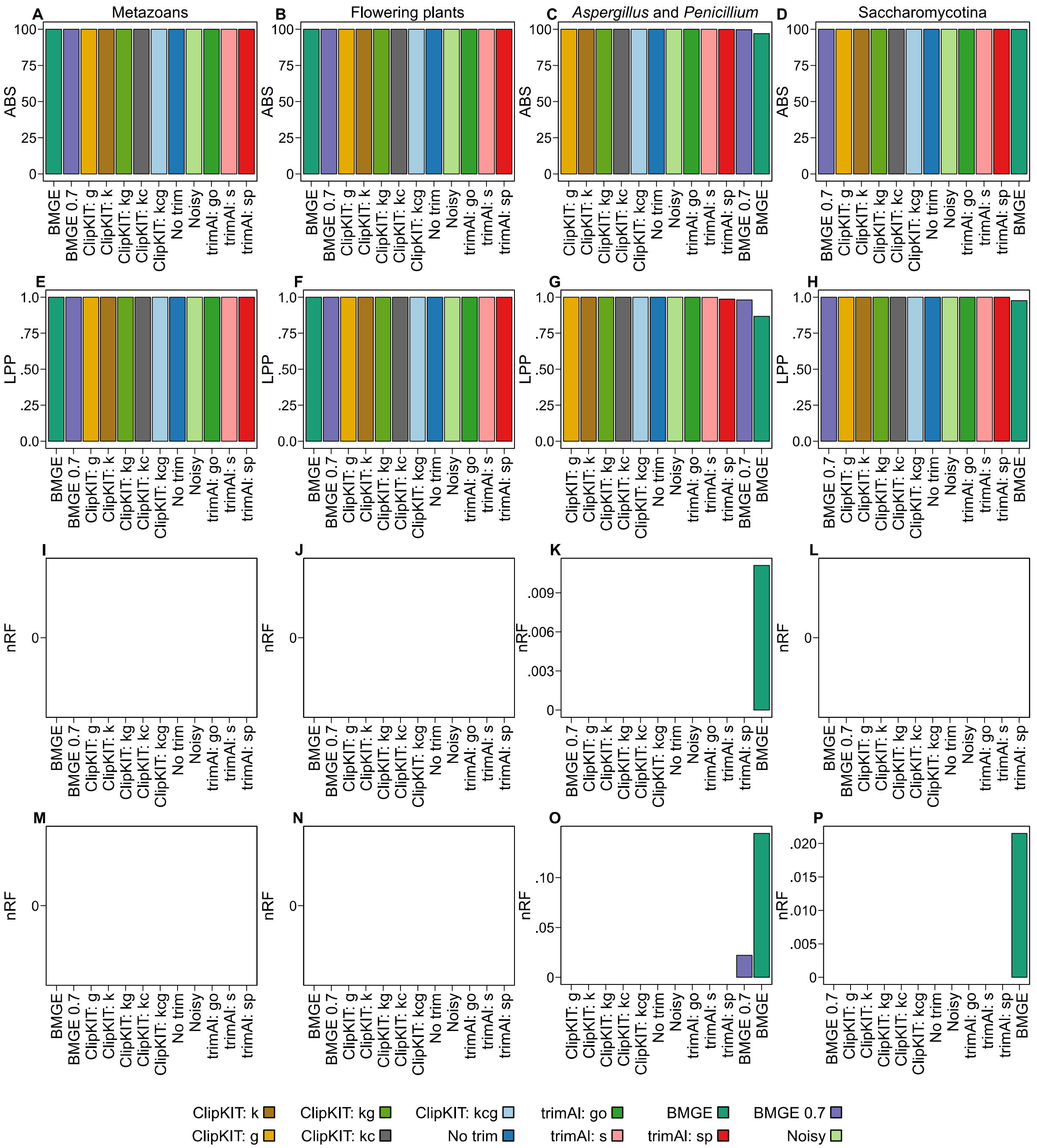


**Supplementary figure 8. Alignment trimming approaches resulted in nearly identical metrics of accuracy and support for species-level inferences using simulated datasets.** Species-level phylogenies were inferred using concatenation and coalescence approaches. Nearly full support was observed for phylogenies inferred using the concatenation (A-D) and coalescence approaches (E-H). Low nRF values indicate the concatenation (I-L) and coalescence approaches (M-P) inferred accurate species-level phylogenies. Concatenation-based metrics are derived from a single tree search.
